# Discovery and pharmacological characterization of nanobodies acting as potent positive allosteric modulators of the calcium-sensing receptor

**DOI:** 10.1101/2024.07.08.602375

**Authors:** Iris Mos, Thomas Zögg, Alexandre Wohlkönig, Anne Mette Helmich Egholm, Sabrina N. Rahman, Els Pardon, Jan Steyaert, Hans Bräuner-Osborne, Jesper M. Mathiesen

**Author notes:** Corresponding author: Jesper M. Mathiesen, Department of Drug Design and Pharmacology, Faculty of Health and Medical Sciences, University of Copenhagen, Denmark.

## Abstract

The calcium-sensing receptor (CaSR) is responsible for sustaining a stable blood calcium concentration. Consequently, genetic and acquired changes in this G protein-coupled receptor can give rise to various calcium homeostasis disorders. Synthetic positive allosteric modulators targeting CaSR are currently used to treat hypercalcemia, but their usage is highly limited due to the high risk of severe hypocalcemia and gastrointestinal intolerance. In this study, we aimed to generate pharmacologically active CaSR-specific nanobodies that could be employed as a new generation of pharmacological tools to investigate the receptor function and potentially serve as a new drug modality for effective treatment of CaSR-related disorders.

Nanobodies were generated by immunization of a llama with CHO cells recombinantly overexpressing a myc-epitope-tagged human CaSR. Following construction of a phage display library representing the repertoire of nanobody genes, nanobodies binding to the CaSR were isolated by FACS of whole HEK293 cells recombinantly overexpressing HA-epitope-tagged human CaSR. Based on sequence comparison, 37 nanobodies from 25 different sequence families were purified and subsequent characterized in vitro for modulation of CaSR signaling. The nanobodies were screened for agonist, as well as positive and negative allosteric modulators activity in *in vitro* cellular assays downstream of CaSR activation. We identified eight pharmacologically active nanobodies acting as positive allosteric modulators that could be divided into five main families based on their sequence identity. The most potent nanobody (Nb4) binding to the extracellular domain of CaSR was slightly more potent than the reference small molecule PAM NPS R-568.

This study describes the discovery and pharmacological characterization of nanobodies acting as potent CaSR positive allosteric modulators. These nanobodies are a new class of pharmacological research tools for the CaSR, which potentially can be developed into new therapeutics in the treatment of CaSR-related disorders.

## Introduction

The calcium-sensing receptor (CaSR) is responsible for maintaining a constant stable serum calcium concentration in the human body (1,2). Accordingly, this class C G protein-coupled receptor (GPCR) is highly expressed in calciotropic tissues (e.g. parathyroid gland, bones and kidney) and is associated with several calcium homeostasis disorders of genetic or acquired origin (3,4). Expressed on the cell surface, the homodimeric CaSR is activated by calcium and further modulated by a number of natural ligands such as other divalent cations, proteinogenic L-amino acids, anions, and polyamines by binding to its large extracellular domain (ECD) (5–7).

Synthetic ligands modulating CaSR activity are of therapeutic importance. The first generation calcimimetic cinacalcet (Sensipar^®^) is a small synthetic molecule acting as a positive allosteric modulator (PAM) of CaSR by binding to the CaSR seven transmembrane domain (7TMD) (8,9). To date, cinacalcet, the first PAM targeting a GPCR to receive market approval, is used as an oral treatment for hypercalcemia in adult patients with parathyroid carcinomas, for primary hyperparathyroidism in patients who cannot undergo parathyroidectomy and for secondary hyperparathyroidism in chronic kidney disease patients receiving hemodialysis (10). Unfortunately, its clinical use is limited due to the high risk of hypocalcemia and gastrointestinal intolerance (11). Over time, the second generation calcimimetic etelcalcetide (Parsabiv^®^), binding to the venus flytrap (VFT) domain of CaSR (9,12) received FDA approval and another small molecule evocalcet (Orkedia_®_,) binding to the same 7TMD binding pocket as cinacalcet (9,13), received approval in Japan as an alternative PAM for cinacalcet in the treatment of secondary hyperparathyroidism in chronic kidney disease patients receiving hemodialysis. Despite improvements, hypocalcemia and gastrointestinal side effects remain and their clinical use is still limited (13–16).

GPCR drug discovery has expanded from synthetic small molecules and peptides towards the generation of GPCR-targeting antibodies or antibody fragments. In contrast to synthetic small molecules, antibodies generally show high target specificity, few off-target effects and offer the opportunity to adjust their *in vivo* half-life by protein engineering (17). In addition to conventional antibodies, camelids and cartilaginous fish express unique heavy chain-only antibodies. The antigen-binding variable heavy domain of a heavy chain only antibody, the VHH domain, has three complementarity determining regions (CDRs) that, especially in the case of CDR3 can adopt long, extended structures in the interaction with their antigen. The VHH fragment, also known as a nanobody, is small in size (∼15 kDa), stable, easy to express and possesses potentially better tissue penetration and faster renal clearance in comparison to the 10-fold larger conventional IgG-based antibody (17). The unique CDR3 structure and the small size allows nanobodies to bind deeply buried cavities such as GPCR binding sites (18) and enzyme active sites (19,20). The therapeutic potential of nanobodies was advanced by the FDA approval of the first nanobody-based therapeutic (caplacizumab) in 2019 for the treatment of acquired thrombotic thrombocytopenic purpura (21).

The present study describes the identification and pharmacological characterization of nanobodies acting as PAMs of the CaSR from a nanobody library obtained through immunization of a llama with whole mammalian cells expressing the human CaSR (hCaSR) followed by whole cell fluorescence-activated cell sorting (FACS)-based nanobody selections and pharmacological screening.

## Materials and Methods

### Materials

Unless stated otherwise, general reagents were purchased from Sigma-Aldrich (St. Louis, MO, USA) and cell culture reagents were bought from Thermo Fisher Scientific (Waltham, MA, USA). The PAM NPS R-568 hydrochloride was purchased from Tocris Bioscience (Bristol, UK), while the negative allosteric modulator (NAM) NPS 2143 hydrochloride was synthesized in-house at the University of Copenhagen, Denmark as previously described (22). Nanobodies were extracted from bacterial cells with TES buffer: 50 mM Tris pH 8.0, 1 mM EDTA, 150 mM NaCl, 20% sucrose (Merck, Darmstadt, DE), 0.4 mg/ml lysozyme, 0.1 mg/ml AEBSF (Carl Roth, Karsruhe, DE) and 1 μg/ml Leupeptin hemisulphate (Carl Roth, Karsruhe, DE). Nanobody selections and pharmacological characterization were performed in assay buffer: Hank’s balanced aalt aolution without Ca^2+^, Mg^2+^ and phenol red (HBSS, Thermo Fisher Scientific, Waltham, MA, USA) supplemented with 20 mM HEPES and adjusted to pH 7.4.

### CaSR plasmid DNA constructs

The human CaSR wild-type (hCaSR WT) sequence (common Q1011 variant of uniprot identifier P41180) was cloned in a pcDNA5 FRT vector to allow stable cell line generation according to the Flp-In technology. The final pcDNA5 FRT hCaSR constructs contained an upstream mGlu5 cleavable signal peptide followed by an N-terminal HA or myc tag. The rat CaSR (rCaSR) WT (uniprot identifier: P48442) construct was made as previously described (23). This construct consists of a metabotropic glutamate receptor subtype 5 (mGlu5) signal peptide and HA tag upstream of the rCaSR WT sequence in a pEGFPN1 background vector.

### Stable cell line generation

The Flp-In technology was applied to establish Flp-In human embryonic kidney 293 (Flp-In HEK293, RRID: CVCL_U421) and Flp-In Chinese hamster ovary (Flp-In CHO, RRID: CVCL_U424) cell lines stably expressing hCaSR WT. At ∼70% confluence, Flp-In HEK293 and Flp-In CHO cells were transfected with a total of 8 μg/dish at a pcDNA5 FRT hCaSR:pOG44 plasmid DNA ratio of 1:9 and 20 μL/dish Lipofectamine 2000. Cells transfected without plasmid DNA were included as negative control. 24h after transfection, the cells were split 1/10, 1/20 and 1/30 into fresh P10 dishes. As from 48h after transfection, the cells were kept under continuous selection pressure with 200 μg/ml hygromycin B for the Flp-In HEK293 cells and 500 μg/ml hygromycin B for the Flp-In CHO cells. Colonies resistant to hygromycin B were pooled together to obtain polyclonal stable Flp-In HEK293 and Flp-In CHO cell lines. A Flp-In HEK293 cell line stably expressing the myc-GABA_A_ δ subunit (previously prepared in a similar fashion) was used in this study as non-CaSR expressing control cell line (24). The GABA_A_ δ subunit is not functional and not present on the cell surface when expressed alone.

### Llama immunization with hCaSR WT recombinantly expressed in Flp-In CHO cells

CaSR-specific nanobodies were generated as previously described (25) with some adjustments. In brief, Flp-In CHO myc-hCaSR WT cells were harvested and stored in aliquots in liquid nitrogen until further use. One llama (Lama glama) was immunized for 6 weeks, by weekly thawing 30-80 million Flp-In CHO myc-hCaSR WT cells, washing cells twice with ice cold HBSS buffer and injecting cells subcutaneously. Four days after the final boost, blood was taken to isolate peripheral blood lymphocytes. From these lymphocytes, RNA was purified and reverse transcribed by RT-PCR to obtain cDNA. The resulting library was cloned into the phage display vector pMESy4 bearing a C-terminal hexa-His tag and a CaptureSelect C-tag sequence (Glu-Pro-Glu-Ala). A nanobody phage display library of 2 billion independent clones was obtained. The phage library was prepared as previously described (25).

### Ethical approval

All animal immunization experiments were executed in strict accordance with good animal practices, following the European Union animal welfare legislation and after approval of the local ethical committee (Committee for the Use of Laboratory Animals at the Vrije Universiteit Brussel). Every effort was made to minimize animal suffering.

### Fluorescence-activated cell sorting (FACS) of whole Flp-In HEK293 cells

For whole cell nanobody selections, mixtures of ∼200,000 freshly cultured Flp-In HEK293 HA-hCaSR WT cells and ∼2 million Flp-In myc-GABA_A_ δ cells were incubated with the phage library (1^st^ round) or rescued phages (2^nd^ round) as described previously (25) and sorted out by FACS on a FACS Aria cytometer (BD Biosciences) in pre-blocked FACS tubes containing 1 ml assay buffer supplemented with 10 mM CaCl_2_ or with 5 μM NPS 2143 and 0.5 mM EDTA. The cells were stained for 45 minutes with 1:1000 PE conjugated anti-HA.11 antibody (BioLegend, #901518, RRID: AB_2629623, San Diego, CA, USA) to visualize CaSR-containing cells and for 5 minutes with 1:1000 TO-PRO-3 iodide (#T3605, Thermo Fisher Scientific, Waltham, MA, USA) to visualize permeable cells. Sort gates were set with 50,000 events as follows: The single cell population was first gated with the forward (FSC) versus side scatter (SSC) plot. Next, the non-permeable cells from the single cell population visualized by negative TO-PRO-3 staining were gated. Finally, the anti-HA PE-positive (i.e. Flp-In HEK293 HA-hCaSR WT) and anti-HA PE-negative (Flp-In HEK293 myc-GABAA δ) cells were sorted from the intact non-permeable single cell population (i.e. non-TO-PRO-3-expressing).

### Nanobody purification

Nanobodies selected for pharmacological characterization were expressed in the periplasm of *E. coli* WK6 cells, extracted and purified by immobilized metal ion affinity chromatography (IMAC) followed by size exclusion chromatography (SEC). First, the pMESy4 construct containing the desired nanobody sequence was transformed into E. coli WK6 competent cells. The pMESy4 construct and bacterial cells were mixed gently and placed on ice for 20 minutes after which the cells were subjected to a heat shock for 1.5 minutes at 42 °C. The heat shocked cells were cooled on ice for 2 minutes and transferred to an Eppendorf tube containing 900 μL LB media. The mixture was shaken at 180 rpm for 1 h at 37 °C, plated on a LB agar plate containing 100 μg/ml ampicillin and 2% glucose and incubated overnight at 37 °C. Second, a pre-culture was prepared in the afternoon by inoculating a single colony in 5 ml LB media containing 100 μg/ml ampicillin and 2% glucose. The pre-culture was grown overnight at 37 °C and 180 rpm. The next morning, 3.5 ml pre-culture was added to 1 L TB media containing 100 μg/ml ampicillin and incubated at 37 °C and 120 rpm for at least 7 hours. The remainder of the pre-culture was mini-prepped and sent for sequencing to confirm the nanobody sequence. At the end of the day, 1 mM isopropyl β-D-1-thiogalactopyranoside (IPTG) was added to the flask to induce nanobody expression overnight at 28 °C and 120 rpm. The induced *E. coli* WK6 competent cells were centrifuged for 10 minutes at 5,000 rpm and the pellet was resuspended in TES buffer to start the lysozyme extraction. The resuspended pellet was supplemented with 0.4 mg/ml lysozyme, 5 mM MgCl_2_ and 1:1000 DNAse I (2500 U/ml, Thermo Fisher Scientific, Waltham, MA, USA) and incubated with head-over-head rotation for 30 minutes at 4 °C. After 30 minutes, the lysed cells were centrifuged for 15 minutes at 20,000 rpm and the nanobody was subsequently purified from the supernatant (i.e. periplasmic extract) via IMAC. IMAC was performed with Ni-NTA resin (GE Healthcare, Chicago, IL, USA) in disposable PD-10 columns. The periplasmic extract was mixed with Ni-NTA resin and incubated for 1 h with head-over-head rotation at 9 rpm prior to addition to an empty column. The Ni-NTA resin was washed four times with 8 ml binding buffer (50 mM Tris pH 8.0, 500 mM NaCl and 10 mM imidazole). Next, the nanobody was eluted in three times each with 700 μL elution buffer (50 mM Tris pH 8.0, 500 mM NaCl and 500 mM imidazole). The nanobody was further purified by SEC using an Enrich SEC 70 10×300 column (BioRad, Hercules, CA, USA). The SEC fractions were analyzed by SDS-PAGE using 4-20% precast protein gels (BioRad, Hercules, CA, USA), pre-stained protein ladder (#26619, Thermo Fisher Scientific, Waltham, MA, USA) and InstantBlue protein staining (Expedeon, San Diego, CA, USA). Pure nanobody fractions were pooled together, concentrated to 175 μM and aliquoted for long term storage at -80 °C.

### Cell culture and seeding protocol for pharmacological characterization

All cell lines utilized in this study were grown at 37 °C and 5% CO2. The HEK293A (RRID: CVCL_6910) and stable Flp-In HEK293 cells were cultured in Dulbecco’s modified Eagle medium (DMEM, #31966021) supplemented with 10% dialyzed fetal bovine serum (dFBS) and 1% 10,000 units/ml penicillin and 10,000 μg/ml streptomycin mixture (pen/strep). The stable Flp-In CHO cells were cultured in F-12K medium (#21127022) supplemented with 10% dFBS and 1% pen/strep. Additionally, the stable Flp-In HEK293 and Flp-In CHO cell lines were cultured in the presence of respectively 200 and 500 μg/ml hygromycin B. For pharmacological characterization, Flp-In HEK293 HA-hCaSR WT cells were seeded in poly-D-lysine-coated clear 96 well tissue culture plates (Corning, Corning, NY, USA) at a density of 50,000 cells/well 18 h prior to assay. Experiments with HA-rCaSR WT required transient transfection of HEK293A cells 24 h prior to assay. HA-rCaSR WT DNA diluted to 30 ng/well DNA in 25 μL/well Opti-MEM was mixed with 0.25 μL/well Lipofectamine 2000 in 25 μL/well Opti-MEM after 5 minutes incubation at room temperature. The DNA:Lipofectamine 2000 mixtures were incubated 20 minutes at room temperature after which 50 μL/well was added to a poly-D-lysine-coated clear 96 well tissue culture plate. Next, 100 μL/well of a 400,000 cells/ml HEK293A cell solution prepared in culture media was added and the plate was placed at 37°C and 5% CO_2_ (final cell density 40,000 cells/well).

### Transient transfection procedure for nanobody characterization at the human mGlu5 receptor

50,000 HEK293 cells/well were transiently transfected in suspension with cDNA encoding pcDNA5 HA-hmGlu5 WT (35 ng/well) and pcDNA5 EAAT3 (glutamate transporter; 15 ng/well) using Lipofectamine 2000 (2.5 μL/well) as transfection reagent. First, cDNA and lipofectamine 2000 solutions were prepared each seperately in OptiMEM, mixed together, followed by an incubation period of 30 minutes at room temperature. After incubation, the cDNA/Lipofectamine2000 solution was added to a single cell suspension in DMEM supplemented with glutaMAX (Gibco; 61965-026), pyruvate, 10% dFBS, + 1% penstrep. Subsequently, 50,000 HEK293 cells/well were added to PDL-coated transparent 96 well plates, and left for overnight incubation at 37°C and 5% CO2. Further characterization of nanobody activity at the hmGlu5 receptor was performed 24 h post transfection using the inositol monophosphate (IP1) accumulation assay (IpOne assay; Cisbio) as described below.

### Inositol monophosphate (IP_1_) accumulation assay

Inositol monophosphate (IP_1_) accumulation was measured using the commercially available HTRF^®^ IP-One accumulation kit (Cisbio Bioassays, Codolet, France) and an EnVision multimode plate reader (PerkinElmer, Waltham, MA, USA). Unless stated otherwise, the cells were washed once with 100 μL/well assay buffer and stimulated for 30 minutes at 37 °C with 50 μL/well ligand(s) diluted in assay buffer supplemented with 20 mM LiCl. Following stimulation, the cells were washed once with 100 μL/well and lysed for 30 minutes at room temperature with 30 μL/well lysis buffer provided with the kit. The lysed cells were diluted 1:2 with assay buffer after which 10 μL/well was pipetted to a white 384-well OptiPlate (PerkinElmer, Waltham, MA, USA). Next, 10 μL of a detection solution consisting of 2.5% IP_1_-d2 conjugate and 2.5% anti-IP_1_ antibody diluted in assay buffer was added to each well. Fluorophore emissions were measured after the plate was stored in the dark for 1 h. The 615/665 nm emission ratios measured upon excitation at 340 nm were converted to IP_1_ concentrations with an IP_1_ standard curve according to the manufacturer’s instructions.

### Advanced Phospho-ERK1/2 assay

Phosphorylation of ERK1/2 at T202 and Y204 was measured with the HTRF^®^ Advanced Phospho-ERK1/2 assay (Cisbio Bioassays, Codolet, France). The cells were handled gently throughout the procedure to avoid agonist-independent ERK1/2 phosphorylation. Cell culture media was aspirated and the cells were washed twice with 50 μL/well Dulbecco’s phosphate buffered saline (DPBS, Thermo Fisher Scientific, Waltham, MA, USA). Next, the cells were pre-incubated for 30 minutes at room temperature with 50 μL/well nanobody or allosteric modulator prepared in assay buffer. After pre-incubation, the liquid was aspirated and the cells were stimulated for 10 minutes at room temperature with 50 μL/well CaCl_2_ solution prepared in assay buffer supplemented with nanobody or allosteric modulator. The agonist solution was removed and the cells were lysed with 50 μL/well lysis buffer (25% lysis solution and 1% blocking reagent diluted in MilliQ) for 1 h at room temperature while shaking at 450 rpm. 16 μL/well lysate was transferred to a white 384-well OptiPlate containing 4 μL/well ERK detection solution (assay buffer supplemented with 2.5% advanced phospho-ERK1/2 d2 antibody and 2.5% advanced phospho-ERK1/2 Eu3+-cryptate antibody) and the plate was incubated for 2 h in the dark. Upon excitation at 340 nm, emission at 615 and 665 nm was measured on an EnVision multimode plate reader (PerkinElmer, Waltham, MA, USA). The resulting 615/665 FRET ratio is used as a direct measure of ERK1/2 phosphorylation.

### Purification of the CaSR extracellular domain and surface plasmon resonance binding assay

The ECD of FLAG-tagged hCaSR was expressed and purified from Sf9 cells as described previously (26). To remove endogenously bound ligands, the purified protein was incubated in 100 mM sodium citrate buffer at pH 5.5 at 4°C overnight. Next, buffer was exchanged to 20 mM HEPES with 150 mM NaCl, concentrated to 8 mg/ml, aliquoted and frozen at -80°C.

Surface plasmon resonance studies were performed on a Biacore X100 by immobilization of an anti-VHH surface MonoRabᵀᴹ Rabbit Anti-Camelid VHH Cocktail (GenScript # A02014-200) to a CM5 chip. Nb4 was captured on the anti-VHH surface and binding kinetics of the diluted purified FLAG-tagged ECD of CaSR were determined in absence or presence of 5 mM Ca^2+^, 5 mM L-Trp or both. The anti-VHH surface was regenerated by 10 mM Glycine pH 2.5.

### Data and Statistical Analysis

The data and statistical analysis comply with the recommendations on experimental design and analysis in pharmacology (27). Data were analyzed with GraphPad Prism v. 8.2 2 and v.10 (GraphPad Software, San Diego, CA, USA). To ensure the reliability of single values, initial pharmacological screening was performed in duplicate and further pharmacological characterization in triplicate. Concentration-response curves were fitted in Prism according to a four-parameter nonlinear regression equation.

## Results

### Generation of CaSR nanobodies using whole recombinant cells

Nanobodies against the human calcium-sensing receptor (hCaSR) were raised and panned using whole cells recombinantly expressing the hCaSR to enable identification of nanobodies targeting the native receptor conformation. The nanobody discovery workflow employed in this study is visually summarized in **Figure 1**. A llama was immunized with whole Flp-In CHO cells recombinantly expressing a myc-tagged wild-type hCaSR (Flp-In CHO myc-hCaSR WT). Four days after the final injection, a phage display library was constructed from isolated peripheral blood lymphocytes by RT-PCR and subcloning of the nanobody cDNA into the pMESy4 phagemid expression vector, allowing generation of a predominantly monovalent nanobody phage library. Nanobody selections were performed using FACS of whole Flp-In HEK293 cells recombinantly expressing HA-tagged hCaSR WT (Flp-In HEK293 HA-hCaSR WT). In an attempt to select for nanobodies that recognize either active or inactive states of CaSR the nanobody phage display selections were performed in the presence of 10 mM CaCl_2_ or in the presence of 5 μM NPS 2143 (NAM) supplemented with 0.5 mM EDTA, respectively. To prevent receptor internalization all selections were performed at 4 °C.

**Figure 1.**
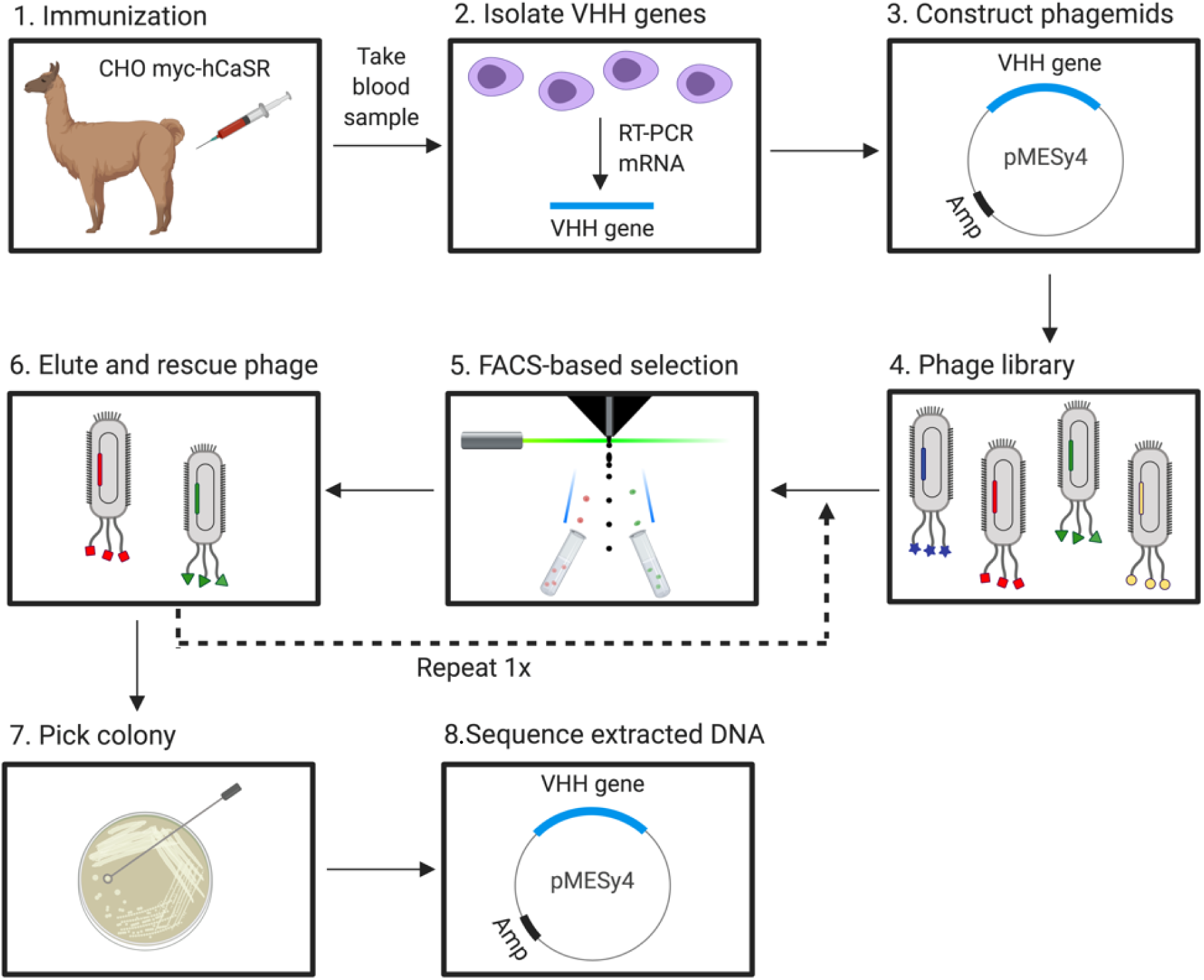
Schematic representation of the CaSR nanobody discovery workflow. A nanobody phage library was created from the isolated nanobody (VHH) genes of a llama immunized with whole Flp-In CHO myc-hCaSR WT cells. Selections of phages were performed in two consecutive rounds each with the following incubation strategies to isolate phages bound to intact Flp-In HEK293 HA-hCaSR WT cells: a) CaSR-expressing cells only, b) Competition incubation strategy - with a 1/10 ratio of CaSR-expressing and non-expressing cells, respectively, c) Pre-incubation strategy - pre-incubation with non-expressing cells for 30 minutes, followed by the competition incubation strategy. Phages bound to intact cells were isolated by FACS, eluted and rescued followed by nanobody DNA extraction and sequencing. Based on sequence analysis, 37 nanobodies were selected for purification and pharmacological characterization. *This image was created with BioRender.com and exported under a paid subscription*.

Isolation of non-specific binders is a common issue in molecular display. To minimize propagation of non-specifically binding nanobodies, different cellular backgrounds and epitopes were used in the immunization and selection steps. All materials (i.e. Eppendorf tubes, FACS tubes, cells and input phages) used were pre-blocked in 1% BSA and 10% FCS for at least 30 minutes to avoid non-specific surface binding of the phage. Typically, whole cell-based phage display selections are performed on antigen-expressing and non-antigen expressing cells in parallel to determine the non-specific cell binders (28). As alternatives and to compare strategies to reduce isolation of non-specific binders in a FACS-based selection procedure three different strategies were employed: 1) incubation with CaSR cells only, where the phage library was incubated for 1.5 hours with only the Flp-In HEK293 HA-hCaSR WT cells, 2) Competition strategy, where the phage library was incubated for 1.5 hours with a 1/10 ratio of Flp-In HEK293 HA-hCaSR WT and non-CaSR expressing Flp-In HEK293 cells (expressing myc-GABA_A_ δ), 3) Preincubation strategy, where the phage library was first pre-incubated for 30 minutes with Flp-In HEK293 myc-GABAA δ cells, the supernatant isolated and then incubated for 1.5 hours with a 1/10 ratio of non-expressing cells, similar to the second strategy. Subsequently for all three strategies, the Flp-In HEK293 HA-hCaSRWT cells (with bound phages) were isolated from non-expressing cells by FACS through direct labelling of the N-terminal HA-tag with a PE-conjugated antibody. The phages were eluted from the isolated cells using trypsin and rescued in TG1 bacterial cells. This selection procedure was repeated in a second round to enrich for CaSR binders, i.e. stabilization of either active or inactive conformation, each with three different strategies to limit isolation of non-specific binders (**Supplemental figure S1**). From each selection strategy, nanobody DNA was extracted from 92 colonies of *E. coli* TG1 infected cells and sanger sequenced. Nanobody DNA sequences obtained after the first and second round were analyzed and grouped in families according to their sequence identity. As expected, certain sequences were found more frequently after the second round of selection indicating an enrichment of these binders. When two or more nanobodies had a high (>80%) CDR3 sequence identity these nanobodies were considered as being within the same family. To exclude nanobody families recognizing background, the sequence alignment was compared with a set of sequences originating from phages binding to non-CaSR expressing (i.e. Flp-In HEK293 myc-GABA_A_ δ cells). Families that re-appeared in these sequencing results were considered non-specific and thus excluded for further characterization. A total of 25 families were identified from sequencing more than 1000 clones (**Supplemental figure S1**). Depending on family size and sequence diversity within a family, we chose 1-3 nanobody clones for purification and pharmacological characterization, resulting in a panel of 37 Nbs for subsequent assaying. Of these 25 families (37 Nbs), 6 families (10 Nbs) originated from the CaCl_2_ condition only, 9 families (14 Nbs) from NPS 2143/EDTA condition only and 10 families (13 Nbs) were found in both conditions (**Supplemental figure S1**).

### Identification of PAM-like CaSR nanobodies and evaluation of selection strategy

The 37 selected nanobodies were purified by IMAC and SEC prior to pharmacological characterization. Each nanobody was initially screened for activity in a functional assay using the IP_1_ accumulation assay (Cisbio IP-One) which is downstream primarily of G_q/11_ activation of CaSR (7,29). Whole HEK293 HA-hCaSR WT cells were pre-incubated for 30 minutes at 37 °C with a fixed nanobody concentration (5 μM) prior to stimulation with the endogenous ligand Ca^2+^. The well-characterized PAM NPS R-568 and NAM NPS 2143 were used as reference compounds. Eight nanobodies (i.e. Nb2, Nb4, Nb5, Nb10, Nb11, Nb15, Nb36 and Nb37) potentiated IP_1_ accumulation in the presence of EC_20_ Ca^2+^ (**Figure 2A**). The PAM effect of these nanobodies, except for Nb5, were recapitulated in the ERK1/2 phosphorylation assay (also predominantly G_q/11_-dependent), when tested in presence of EC_20_ Ca^2+^ (**Figure 2B)**. No nanobodies with agonist or significant NAM activity were identified in neither IP_1_ accumulation or ERK1/2 phosphorylation assay when all nanobodies were screened in the absence or in the presence of EC_80_ Ca^2+^, respectively (**Supplemental Figure S2**).

**Figure 2.**
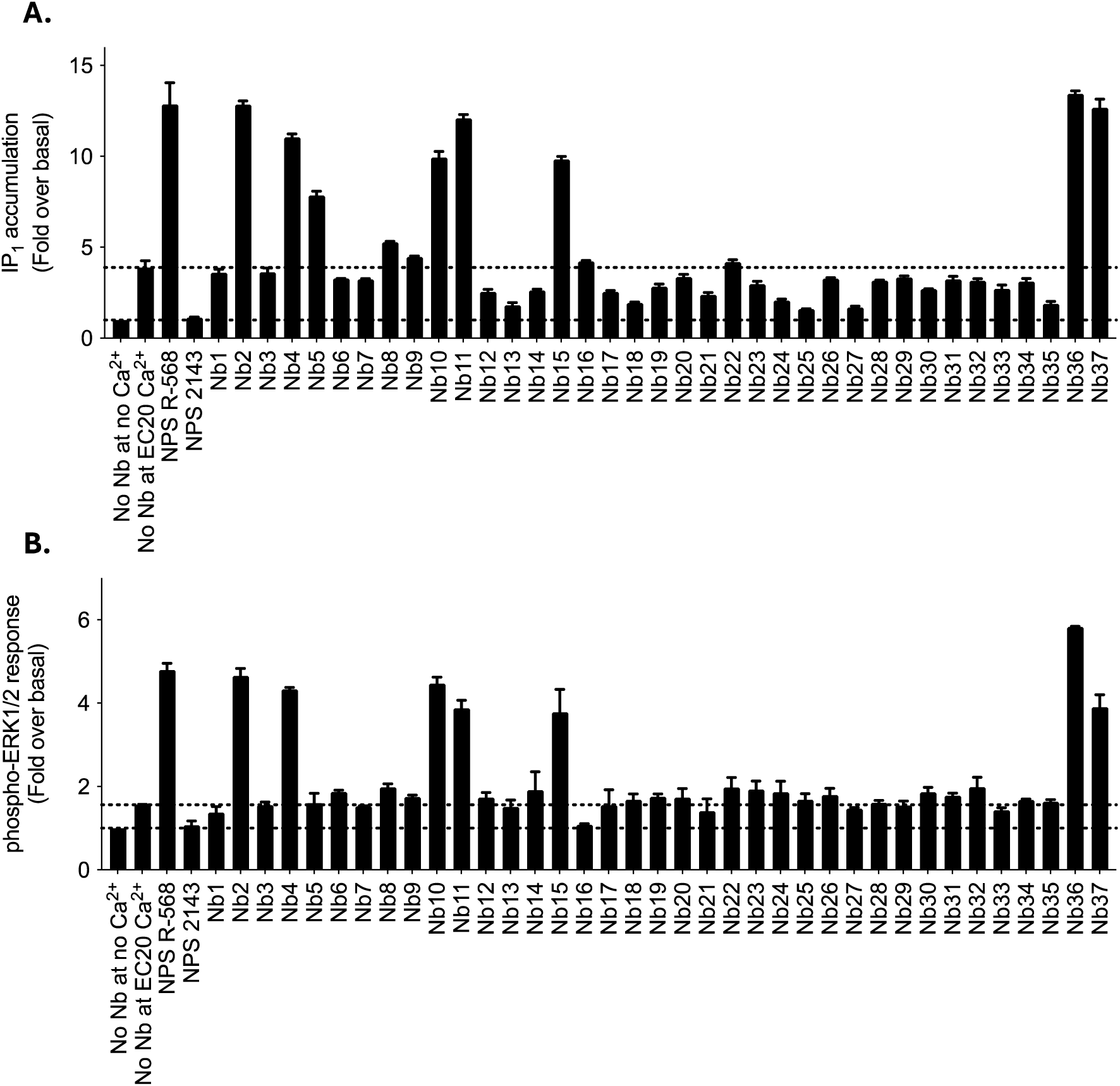
Pharmacological characterization of 37 selected CaSR nanobodies library for PAM activity in a Flp-In HEK293 cell line recombinantly expressing the HA-tagged human CaSR WT. Testing of purified nanobodies at 5 µM for PAM activity at EC_20_ Ca^2+^ in (A) the IP-One accumulation and (B) the phospho-ERK1/2 assays. Data are presented as mean ± S.D. of a single experiment performed in duplicate.

Testing of the eight pharmacologically active nanobodies in a concentration-dependent manner for potentiation of an EC_20_ Ca^2+^ in the IP_1_ accumulation assay showed a potency rank order of Nb4 > NPS R-568 > Nb36 >Nb2 > Nb15 > Nb10 > Nb11 > Nb37 >>> Nb5 (**Figure 3A**, **Table 1**) for the HA-tagged hCaSR. In line with the high sequence identity of the human and rat ortholog of CaSR (93.6% for full-length and 95.7% for the ECDs), the nanobodies showed comparable rank order and potency when tested on the rat CaSR (rCaSR) (**Figure 3B**, **table 1**), enabling further characterization in *in vivo* rat models.

**Figure 3.**
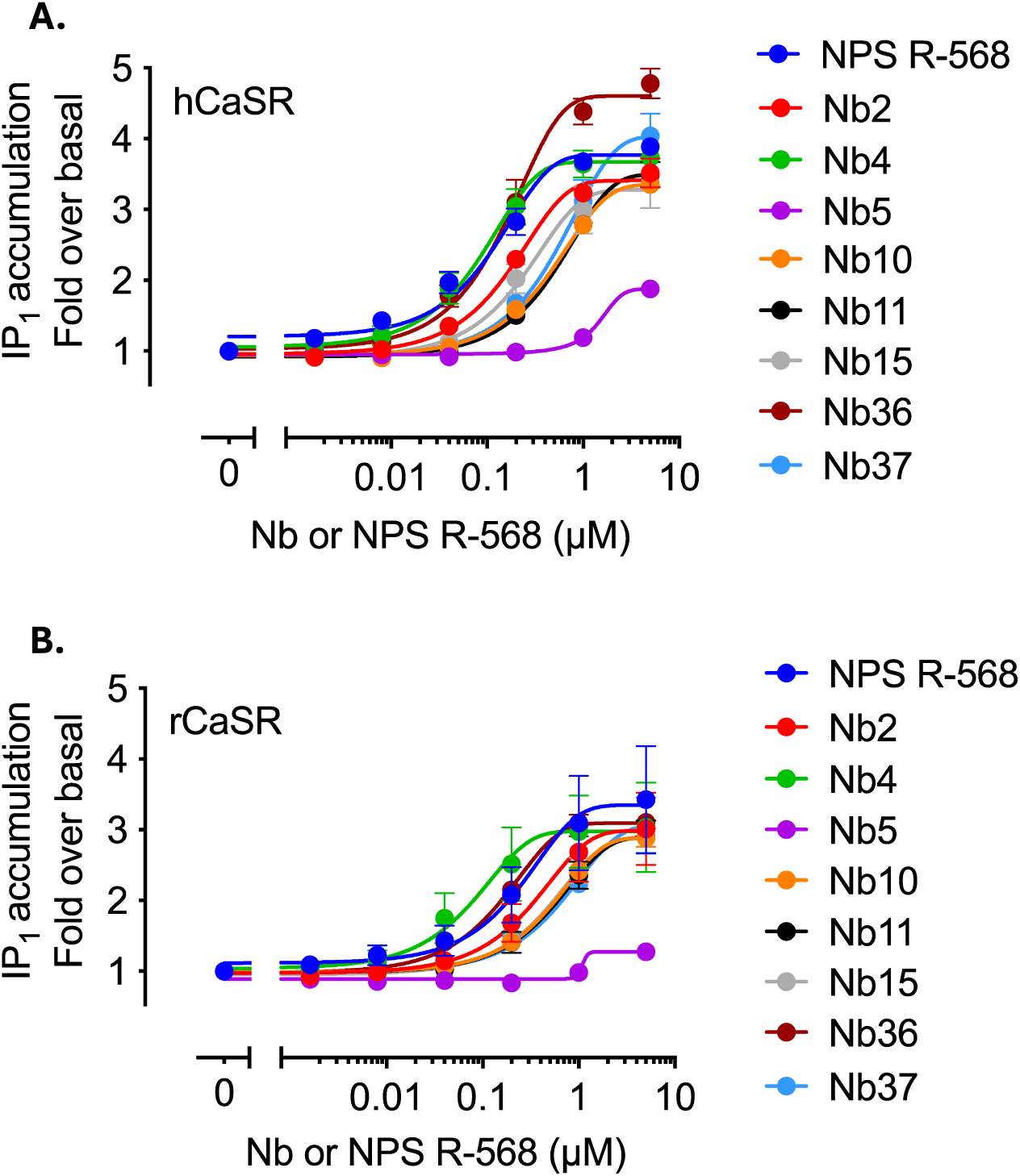
Nanobody and NPS R-568 concentration-response curves at EC_20_ Ca^2+^ for HA-tagged CaSR human and rat ortholog. IP_1_ accumulation of (A) HA-tagged human (hCaSR) or (B) HA-tagged rat CaSR (rCaSR) upon stimulation with EC_20_ Ca^2+^ and increasing nanobody or NPS R-568 concentrations. Data are presented as mean ± S.E.M. of three independent experiments performed in triplicate.

**Table 1.** Nanobody potency on human and rat CaSR. Potency values as pEC_50_ and maximal response (max. response) as fold over basal of the nanobodies and small molecule PAM NPS R-568 obtained for human and rat HA-tagged CaSR (from curves in Figure 3). N.D., not determined.

|  | Human ortholog |  | Rat ortholog |  |
| --- | --- | --- | --- | --- |
|  | Potency<br>(pEC<sub>50</sub> ± S.E.M.) | Max. response<br>(fold over basal ±<br>S.E.M.) | Potency<br>(pEC<sub>50</sub> ± S.E.M.) | Max. response<br>(fold over basal ±<br>S.E.M.) |
| **NPS R-568** | 6.92 ± 0.15 | 3.89 ± 0.07 | 6.59 ± 0.22 | 3.43 ± 0.76 |
| **Nb2** | 6.71 ± 0.08 | 3.52 ± 0.21 | 6.46 ± 0.07 | 3.01 ± 0.51 |
| **Nb4** | 7.08 ± 0.13 | 3.74 ± 0.08 | 7.09 ± 0.09 | 3.03 ± 0.63 |
| **Nb5** | N.D.* | 1.88 ± 0.05 | N.D.* | 1.28 ± 0.12 |
| **Nb10** | 6.33 ± 0.03 | 3.36 ± 0.09 | 6.29 ± 0.02 | 2.89 ± 0.13 |
| **Nb11** | 6.28 ± 0.07 | 3.50 ± 0.17 | 6.24 ± 0.11 | 2.90 ± 0.23 |
| **Nb15** | 6.59 ± 0.03 | 3.38 ± 0.36 | 6.31 ± 0.07 | 2.88 ± 0.13 |
| **Nb36** | 6.78 ± 0.17 | 4.78 ± 0.21 | 6.78 ± 0.03 | 3.10 ± 0.12 |
| **Nb37** | 6.22 ± 0.05 | 4.04 ± 0.32 | 6.00 ± 0.28 | 3.07 ± 0.39 |

\*Greater than the highest tested concentration.

An examination of the origin of the eight pharmacologically active nanobodies in relation to the selection strategies showed that all PAM nanobodies were obtained with the CaCl_2_ condition and none in the NPS 2143/EDTA condition. The active nanobodies belonged to five distinct nanobody families since Nb4 was highly similar to Nb11 and Nb36 (89.9% and 90.6% sequence identity, respectively) and Nb10 was highly similar to Nb37 (97.8 % sequence identity). The total number of nanobody sequences identified with the CaCl_2_ condition within each family were evaluated in relation to the selection strategy and round (**Table 2**). In general, most nanobodies from the active families were observed after round 2, in agreement with an enrichment over selection rounds. The CaSR cell only strategy appeared less effective for enrichment. Interestingly, particular families appeared to be mostly or solely obtained with one of the selection strategies.

**Table 2.** Pharmacologically active nanobody summary table. Schematic overview of the distribution into families of the pharmacologically active nanobodies together with the total number of sequences/clones identified within each family and the conditions under which these clones were identified. Conditions where no clones within a family were observed are indicated with ‘no’.

| Pharmacologically active nanobodies distributed into families | Number of sequences/clones identified from this family | CaSR cells only |  | Competition with non-CaSR expr. cells |  | Pre-incubation and competition with non-CaSR expr. cells |  |
| --- | --- | --- | --- | --- | --- | --- | --- |
|  |  | R1 | R2 | R1 | R2 | R1 | R2 |
| Fam 1: Nb2 | 8 | no | no | no | no | no | 8 |
| Fam 2: Nb4+Nb11+Nb36 | 18 | 1 | no | no | 2 | no | 15 |
| Fam 3: Nb5 | 4 | no | no | no | 1 | 2 | 1 |
| Fam 4: Nb10+Nb37 | 7 | no | no | no | 6 | no | 1 |
| Fam 5: Nb15 | 4 | no | 2 | no | 1 | no | 1 |

### Detailed pharmacological characterization of CaSR nanobodies

Further pharmacological characterization of the PAM nanobodies was performed with at least one representative from each family of pharmacologically active PAM nanobodies, except for Nb2 which was not characterized further due to low yield upon expression and purification. Also, since Nb15 did not have a stop codon at the common stop-codon position of nanobodies, the Nb15 was substituted for Nb45 in the subsequent characterization. Nb45 has an identical sequence as Nb15 and a stop-codon at the expected position. As similar maximal PAM responses were observed with and without a 30 minutes pre-incubation step in the IP1 accumulation assay (**Supplementary Figure S3**), further characterization was performed without a 30 min pre-incubation step.

To provide a more thorough pharmacological characterization, the selected nanobodies were characterized in an agonist potency shift assay format. Each nanobody and NPS R-568 as a positive control were tested for their ability to left-shift (i.e. potentiate) the Ca^2+^ concentration-response curve (CRC) (**Figure 4**). The potency shift plots confirmed the poor PAM properties of Nb5, due to almost no leftward shift of the Ca^2+^ CRC, whereas the other nanobodies all left-shifted the CRC of Ca^2+^ in a concentration-dependent manner. Nb4 appeared to more potently left shift the Ca^2+^ CRC as compared to the small molecule PAM NPS R-568. Interestingly, nanobodies appeared to affect the Hill slope of the Ca^2+^ differentially. In addition, some nanobodies slightly decreased the maximal response (E_max_) of Ca^2+^, while increasing the potency of Ca^2+^ (eg. Nb36). These different functional profiles could indicate different binding modes of the nanobodies to CaSR, thereby affecting the receptor dynamics differently.

**Figure 4.**
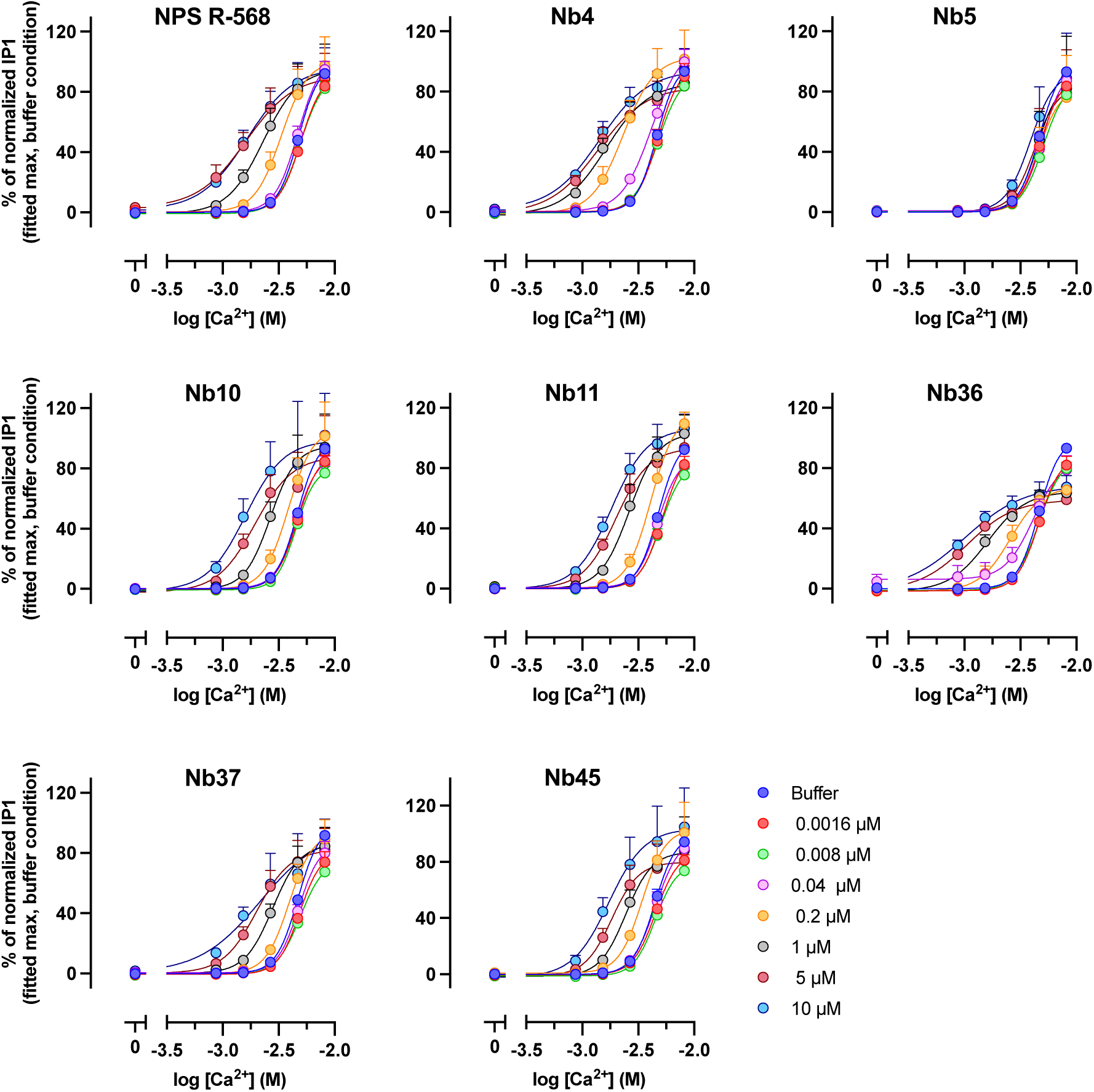
PAM nanobodies’ ability to increase the potency of Ca^2+^ at the HA-tagged human CaSR. Flp-In HEK293 HA-hCaSR WT cells were tested in the IP_1_ accumulation assay using increasing concentration of nanobodies in the presence of Ca^2+^ at different concentrations. The data are shown as percentage of the fitted max when Ca^2+^ was tested in absence of nanobody. Nb2 was not tested in the agonist potency shift plot as it was very poorly expressed. Data are given as means ± S.E.M of three independent experiments performed in triplicate.

In general, nanobodies possess good target selectivity. To validate selectivity towards CaSR, the selected nanobodies were tested for activity at the human mGlu5, a closely related family C GPCR, for which PAM-like nanobodies have previously been described (30). When tested at 10 µM, none of the nanobodies affected the potency of mGlu5 agonist quisqualic acid, whereas the mGlu5 PAM CDPPB clearly shifted the quisqualic acid CRC to the left (**Figure 5A**). These data further corroborate that the observed PAM responses at CaSR likely are due to specific activity at the CaSR.

**Figure 5.**
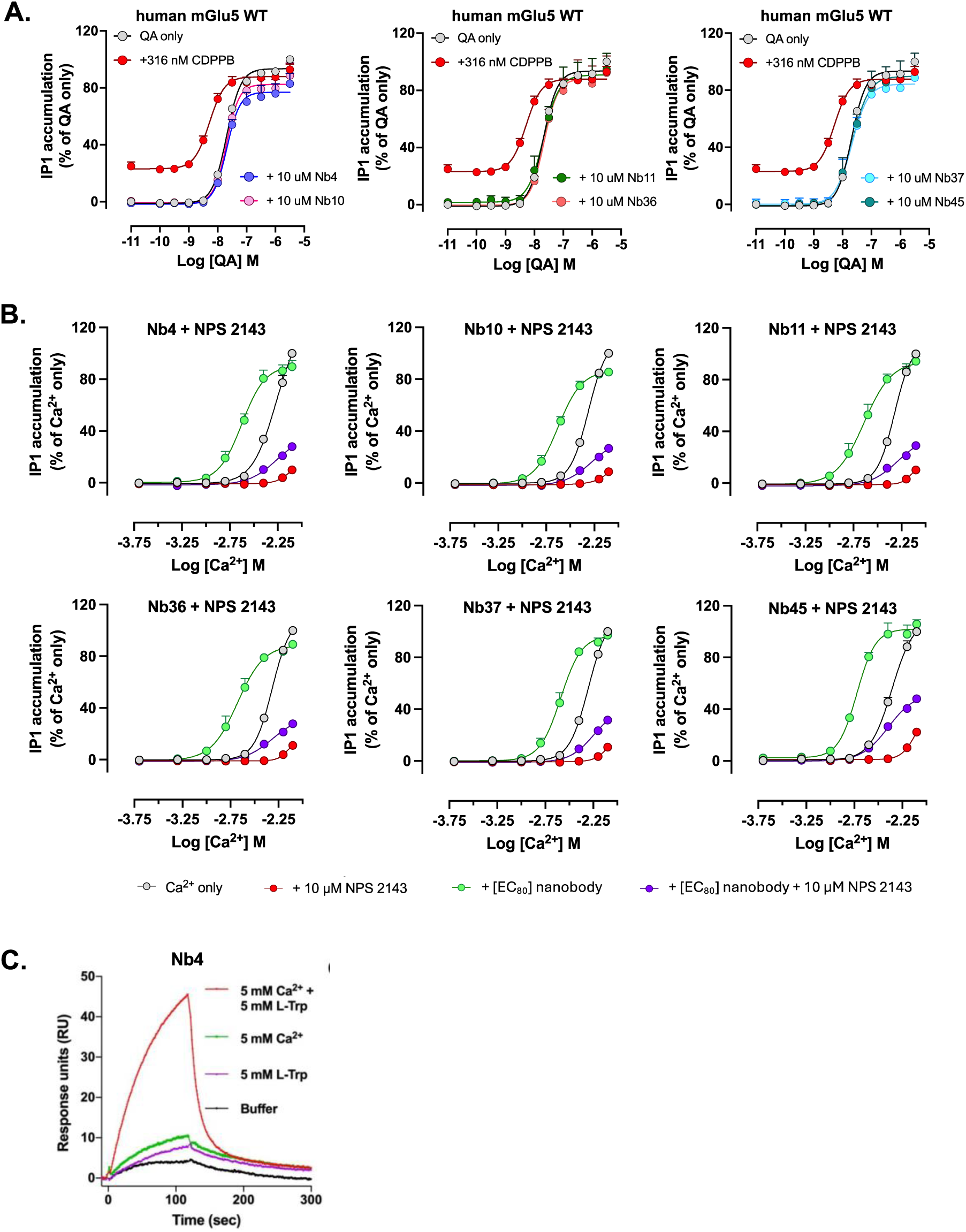
Nanobody PAM selectivity and dependence on an active CaSR conformation. (A) A mGlu5 PAM (CDPPB), but not the nanobodies, affect activation of the HA-tagged human mGlu5 WT (HA-mGlu5 WT). (B) the CaSR NAM NPS 2143 diminish nanobody potentiation of the Ca^2+^ response at HA-tagged CaSR WT. (C) Nb4 only binds efficiently to the purified CaSR ECD when in presence of both Ca^2+^ and L-Trp. Data in (A) and (B) are means ± S.E.M of three independent experiments performed in triplicate. Means are normalized to the max response by (A) the mGlu5 agonist quisqualic acid alone (QA alone) or (B) Ca^2+^ alone. Data in (C) are from one representative study and given as response units (RU), reflecting an increase in mass by binding of the CaSR ECD to Nb4 immobilized on the chip surface.

To induce G protein-mediated signaling, the CaSR undergoes conformational changes both in the VFT and the 7TMD upon agonist binding (7,31). It is well established that 7TMD-binding NAMs such as NPS 2143 inhibit agonist-induced signaling of the CaSR by stabilizing the 7TMD in an inactive conformation (9,32,33). In presence of NPS 2143, the nanobody-induced potentiation of Ca^2+^-mediated signaling was blocked albeit not to the same extent as by Ca^2+^ alone, again indicating a potent but quenchable PAM effect of the nanobodies (**Figure 5B**).

Finally, to provide an indication of the binding site for one of the most potent nanobodies and explore PAM-dependence on the presence of L-Trp and Ca^2+^ for binding to the CaSR, Nb4 was tested for binding to a purified ECD of the CaSR by surface plasmon resonance. L-Trp, which is typically found in the common amino acid binding pocket of ECD, was removed by lowering the pH during purification as previously described (26). Following immobilization of Nb4 to the surface, the CaSR ECD in the absence or presence of Ca^2+^ or L-Trp or both was flowed over the Nb4-coated surface. Interestingly, the strongest binding was observed only in presence of both L-Trp and Ca^2+^, indicating that Nb4 acts as a PAM by binding to an active-like conformation of the CaSR ECD (**Figure 5C**).

## Discussion and Conclusions

In the present study, we successfully raised a llama-derived CaSR nanobody library using CaSR overexpressing whole cells for immunization and selection. Pharmacological characterization led to the discovery of eight nanobodies acting as potent CaSR PAMs, of which 5 belong to different sequence families.

By performing nanobody selections on whole cells, we envisioned that we would be able to induce and stabilize different native CaSR conformations by adding ligands during selection. We attempted to induce active- and inactive-like CaSR conformations by addition of 10 mM CaCl_2_ or 5 μM NPS 2143 supplemented with 0.5 mM EDTA, respectively, during panning. In agreement with their pharmacological profiles, the eight PAM-acting nanobodies were all solely obtained from the 10 mM CaCl_2_ selection. However, of all nanobodies tested, no potent nanobodies with NAM activity were identified for the 5 μM NPS 2143 / 0.5 mM EDTA condition. As mentioned, CaSR can be activated by a wide range of natural ligands, such as Ca^2+^ and polyamines. These ligands are also naturally present in the blood stream of the llama during immunization, thereby potentially shifting the conformation equilibrium towards an active conformation during immunization with whole Flp-In CHO myc-hCaSR WT cells. Further stabilization of the ECD in an inactive conformation also during selection, eg. by more a strict control of Ca^2+^ and polyamines, may be required to isolate NAM-like nanobodies using the whole cell selection approach.

In line with the enrichment of binders in molecular displays by performing several selection rounds, most PAM nanobody families first emerged after the second selection round. A deselection approach by addition of and/or preincubation with non-CaSR expressing cells to remove non-specific binders appeared to be key for isolating the CaSR PAM nanobodies (**Supplemental figure S1**). Interestingly, some families were mostly or solely obtained with one of the selection strategies employed. This observation emphasizes the importance of employing different selection strategies in parallel, at least in the current study.

We were able to confirm the PAM activity of the most potent nanobodies in the orthogonal phosphor-ERK1/2 assay. Only the PAM activity of Nb5 could not be confirmed which aligns with it displaying the weakest PAM in the IP_1_ assay (**Figure 2**). Where the phospho-ERK1/2 assay measures a natural cell response (i.e. phosphorylation of ERK1/2), the IP-One accumulation assay measures non-natural accumulation of the second messenger IP_1_. This difference could make the IP_1_ accumulation assay more sensitive to picking up activity from low potent ligands such as Nb5. Although the nanobodies had no effect in the mGlu5 assay, future in vitro studies should be performed to confirm selectivity. Even though immunization and selection were performed using different N-terminally tagged CaSR to avoid isolation of binders to the tags, PAM activity of the nanobodies at the untagged CaSR should also be tested.

The pharmacological characterization indicates a PAM-like mechanism of action of the nanobodies. When applied alone, the pharmacologically active nanobodies and reference PAM NPS R-568 did not induce IP_1_ accumulation or ERK1/2 phosphorylation in the absence of added Ca^2+^, and increased potency of Ca^2+^except for the low potency Nb5. For Nb4 that displayed high PAM potency, we found that the nanobody bind the ECD of CaSR in a Ca^2+^ and L-Trp dependent manner. Together this indicates that Nb4 recognizes and stabilizes a fully active closed-closed state of the CaSR. ECD-binding of the remaining nanobodies is yet to be tested.

Nanobodies have proven useful as tools in GPCR research. They can be used to visualize the GPCR in native cells or tissues, unveil novel potential drug targeting sites and act as chaperones in GPCR crystallization (18,34). The first full-length structure of a class C metabotropic glutamate receptor subtype 5 homodimer was resolved in presence of a PAM nanobody (30). Although it does not appear necessary to stabilize the CaSR by nanobodies to obtain active and inactive cryoEM structures (7–9,31,32), an inactive CaSR with a NAM-like nanobody binding to the ECD was recently reported (33). The nanobodies described in this study may offer the opportunity to facilitate determination of a higher resolution, active conformation, full-length CaSR structure.

One drawback of current CaSR-targeting PAMs is the high risk for severe hypocalcemia and gastro-intestinal adverse effects (11,14,15). Depending on the mechanism of adverse effects of the current CaSR drugs, the CaSR nanobody modality could have a potential for lowering adverse effects. CaSR is a widely expressed GPCR and thus adverse effects through activation of CaSR in different tissues such as the gastrointestinal tract is not surprising (29). As a larger drug modality and with lower lipophilicity than small molecule PAMs, nanobodies could potentially overcome some of these adverse effects although etelcalcetide, a highly charged peptide PAM for CaSR, does not show a reduced side effect profile (14,35).

In conclusion, a CaSR nanobody library was generated leading to the discovery of several nanobodies acting as CaSR PAMs for the first time. Future research will focus on resolving the binding mechanism and sites of the CaSR PAM nanobodies and further explore the therapeutic potential of these nanobodies in *in vivo* CaSR-related disease models.

## Acknowledgements

We thank Eva Beke for technical assistance during nanobody discovery and Dr. Simon R. Forster for fruitful discussions and assistance with fundraising. This work benefited from access to the Nanobodies4Instruct center and acknowledge the use of resources of Instruct-ERIC, part of the European Strategy Forum on Research Infrastructures (ESFRI). We acknowledge the Research Foundation Flanders (FWO) for their support of the nanobody discovery.

This project has received funding from the European Union’s Horizon 2020 research and innovation programme under grant agreement NO 675228, Fonden til Lægevidenskabens Fremme and by the INSTRUCT European Research Infrastructure Consortium.

## Abbreviations

7TMD: seven transmembrane domain
CaSR: calcium-sensing receptor
CDR: complementarity-determining regions
CHO: Chinese hamster ovary
CRC: concentration-response curve
dFBS: dialyzed fetal bovine serum
DMEM: Dulbecco’s modified Eagle medium
DPBS: Dulbecco’s phosphate buffered saline
ECD: extracellular domain
ELISA: enzyme-linked immunosorbent assay
ERK1/2: extracellular signal-regulated kinase 1 and 2
FACS: fluorescence-activated cell sorting
GPCR: G Protein-coupled receptor
HBSS: Hank’s balanced salt solution
HEK: human embryonic kidney
HTRF: homogeneous time resolved fluorescence
IMAC: immobilized metal ion affinity chromatography
IP_1_: inositol monophosphate
IPTG: isopropyl β-D-1-thiogalactopyranoside
mGlu5: metabotropic glutamate receptor subtype 5
NAM: negative allosteric modulator
PAM: positive allosteric modulator
SDS-PAGE: sodium dodecyl sulphate-polyacrylamide gel electrophoresis
SEC: size exclusion chromatography
VFT: venus flytrap
VHH: variable heavy domain of a heavy chain-only antibody.

## Supplementary

**Supplemental figure S1.**
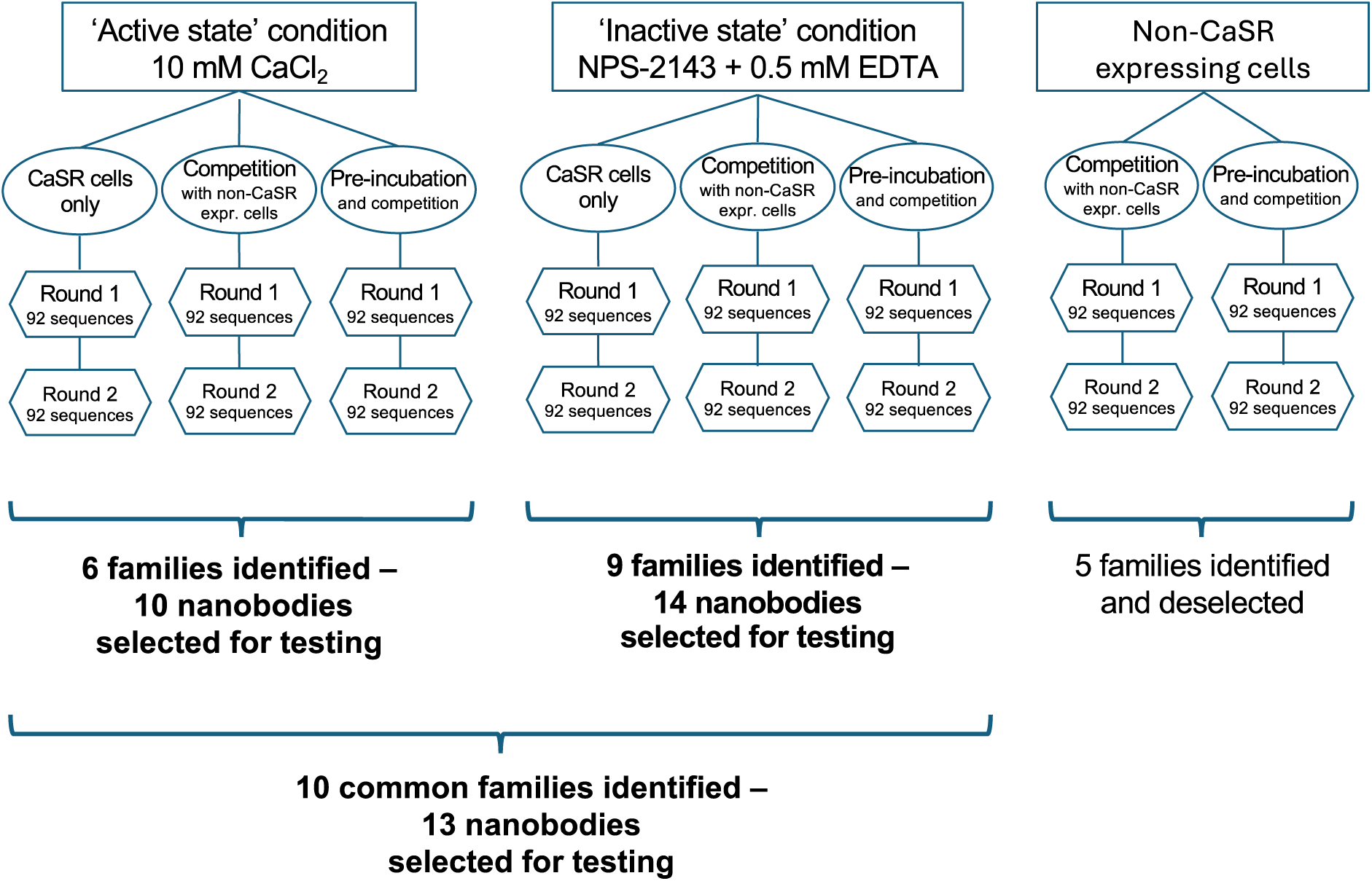
Selection strategies and origin of nanobodies chosen for pharmacological characterization. Two different selection conditions (active state condition and inactive state condition) were used for CaSR-expressing cells along with non-CaSR expressing cells to identify non-specific binders. For the active and inactive state conditions, phage selections were performed in 2 consecutive rounds each with the following incubation strategies to isolate phages bound to intact Flp-In HEK293 HA-hCaSR WT cells by FACS: a) CaSR-expressing cells only, b) Competition incubation strategy - with a 1/10 ratio of CaSR-expressing and non-expressing cells, respectively, c) Pre-incubation strategy - pre-incubation with non-expressing cells for 30 minutes, followed by the competition incubation strategy. Following sequence analysis 25 different families were identified. Six families were identified only in the CaCl_2_ condition, nine families were identified only in the NPS 2143/EDTA condition and 10 families were identified in both conditions. To exclude nanobody families recognizing background, the sequence alignment was compared with a set of sequences originating from phages binding to non-CaSR expressing Flp-In HEK293 myc-GABAA δ cells). Five families that re-appeared in the sequencing results of non-expressing CaSR cells were considered non-specific and excluded.

**Supplemental figure S2.**
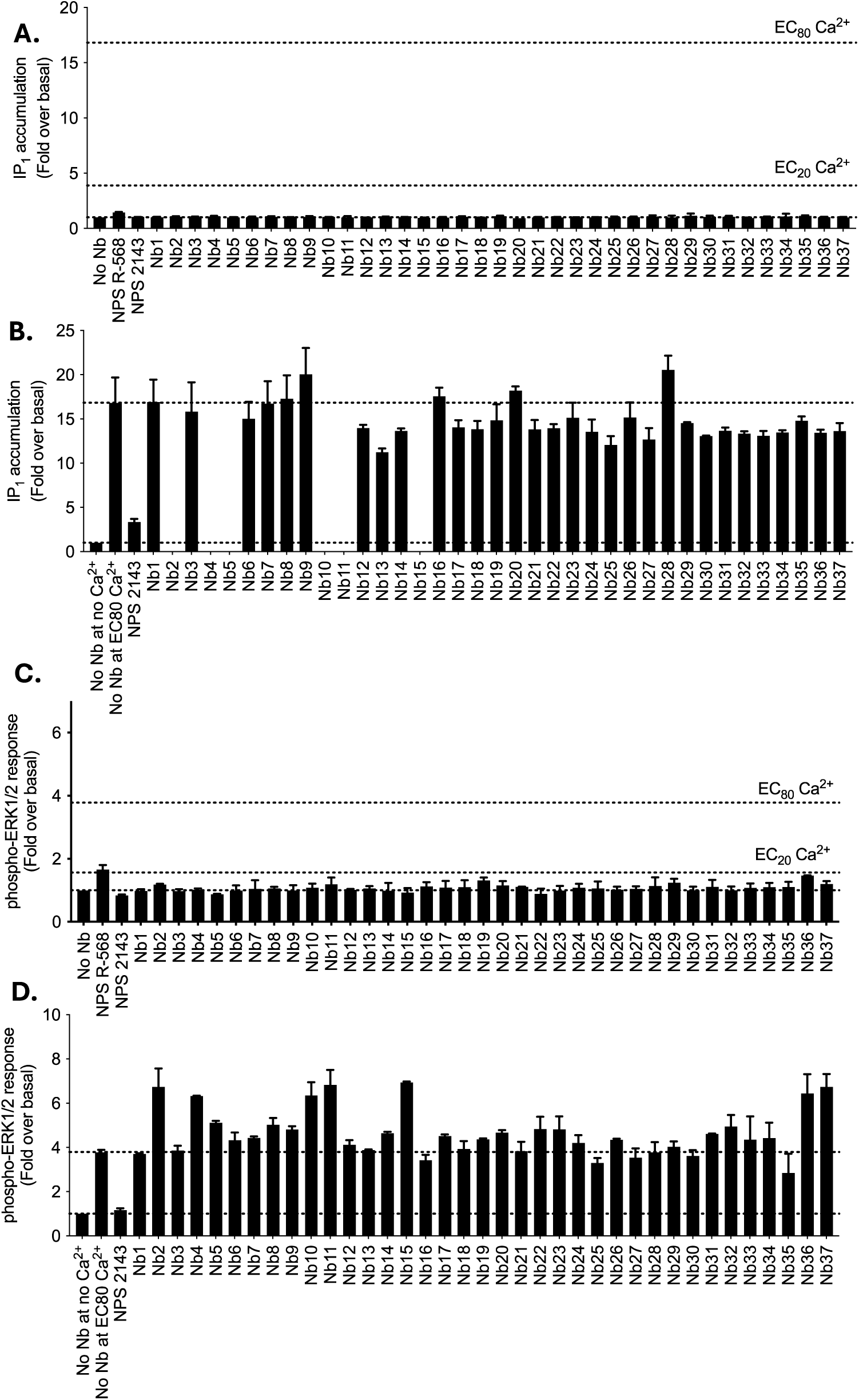
Pharmacological characterization of 37 selected CaSR nanobodies library for agonist and NAM activity in a Flp-In HEK293 cell line recombinantly expressing the HA-tagged human CaSR WT. Testing of purified nanobodies at 5 µM for (A) agonist activity in the absence of Ca^2+^ and (B) NAM activity at EC_80_ Ca^2+^ in the IP-One accumulation assays, and (C) agonist activity in the absence of Ca^2+^ and (D) NAM activity at EC_80_ Ca^2+^ in the phospho-ERK1/2 assays. Data are presented as mean ± S.D. of a single experiment performed in duplicate.

**Supplemental figure S3.**
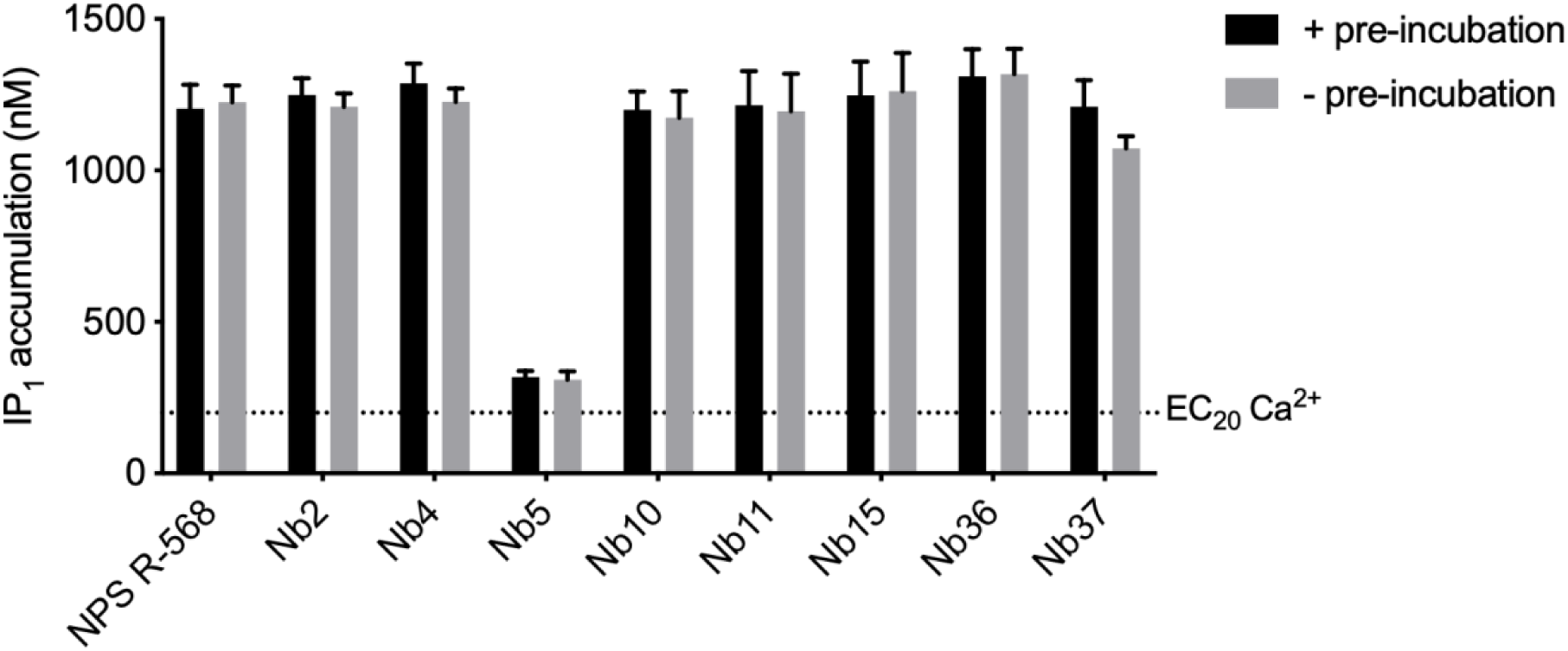
Nanobody and NPS R-568 pre-incubation test. IP_1_ accumulation response obtained for 5 μM NPS R-568 or pharmacologically active nanobodies with (+) or without (-) 30 minutes pre-incubation at 37 °C prior to stimulation with EC_20_ Ca^2+^. The dotted line indicates the IP_1_ response at EC_20_ Ca^2+^ only. Data are mean ± S.E.M. of three independent experiments performed in triplicate.

